# A multi-gene region targeted capture approach to detect plant DNA in environmental samples: A case study from coastal environments

**DOI:** 10.1101/2021.07.03.450983

**Authors:** Nicole R. Foster, Kor-jent van Dijk, Ed Biffin, Jennifer M. Young, Vicki Thomson, Bronwyn M. Gillanders, Alice Jones, Michelle Waycott

**Affiliations:** The University of Adelaide; State Herbarium of South Australia; Flinders University; University of Adelaide; State Herbarium of South Australia and The University of Adelaide

## Abstract

Metabarcoding of plant DNA recovered from environmental samples, termed environmental DNA (eDNA), has been used to detect invasive species, track biodiversity changes and reconstruct past ecosystems. The P6 loop of the *trnL* intron is the most widely utilized gene region for metabarcoding plants due to the short fragment length and subsequent ease of recovery from degraded DNA, which is characteristic of environmental samples. However, the taxonomic resolution for this gene region is limited, often precluding species level identification. Additionally, targeting gene regions using universal primers can bias results as some taxa will amplify more effectively than others. To increase the ability of DNA metabarcoding to better resolve flowering plant species (angiosperms) within environmental samples, and reduce bias in amplification, we developed a multi-gene targeted capture method that simultaneously targets 20 chloroplast gene regions in a single assay across all flowering plant species. Using this approach, we effectively recovered multiple chloroplast gene regions for three species within artificial DNA mixtures down to 0.001 ng/µL of DNA. We tested the detection level of this approach, successfully recovering target genes for 10 flowering plant species. Finally, we applied this approach to sediment samples containing unknown compositions of environmental DNA and confidently detected plant species that were later verified with observation data. Targeting multiple chloroplast gene regions in environmental samples enabled species-level information to be recovered from complex DNA mixtures. Thus, the method developed here, confers an improved level of data on community composition, which can be used to better understand flowering plant assemblages in environmental samples.

## 1 Introduction

Environmental DNA (eDNA) is a rapidly growing field of research and has been applied extensively to monitor site-based vegetation change over periods of hundreds to thousands of years using samples from soil cores (Willerslev et al., 2003; Willerslev et al., 2014; Parducci et al., 2013; Zimmermann et al., 2017; Del Carmen Gomez Cabrera et al., 2019). As plants are sedentary, they are a reliable reflection of their environment at a specific point in time (Yoccoz et al., 2012), therefore, reconstruction of plant communities through time can represent environmental conditions and how they have changed at the sampled location. Such information can be used to predict future changes, inform management (Fordham et al., 2016; Balint et al., 2018; Brown & Blois, 2001) and provide insight to uncover species extinctions and/or introductions, elucidate past climate conditions and assess ecosystem trajectories (Thomsen & Willerslev, 2015; Ruppert, Kline & Rahman, 2019). In addition, changes in plant community composition can shed light on historical human impacts and agricultural practices (Giguet-Covex et al., 2014; Pansu et al., 2015).

The ability to take an environmental sample and accurately determine the plant community composition contained within, as remnant fragments of tissue or DNA, is most commonly achieved through the process of DNA metabarcoding. Metabarcoding involves Polymerase Chain Reaction (PCR) amplification of a short, but highly variable, gene region that is flanked by conserved regions. The design of PCR amplification primers target these conserved regions, which allows amplification of the variable region across all plant taxa present in an environmental sample (Murchie et al., 2020). The variable region that is often used to recover plant DNA in environmental samples is the P6 loop of the chloroplast *trnL* (UAA) intron (Taberlet et al., 2006). This gene region was adopted due to being short enough to amplify DNA in environmental samples (10–143 bp), whilst still possessing enough discriminatory information to distinguish many groups of plant taxa. Unfortunately, reliance on a single, short, gene region creates limitations to generating accurate results, as species can only be detected if this particular region remains present in the genomic DNA extracted, and if the primer binding sites are intact to allow successful amplification. Additionally, targeting only one gene region using a single pair of universal primers creates bias in the results as primers may preferentially bind to certain taxa, creating an unreliable representation of the vegetation community (Pedersen et al., 2015). Further, recovering the short section of the *trnL* gene region does not ensure taxonomically high-resolution results, given the lowered discrimination ability of this region due to the short length. Other commonly used plant barcodes such as *rbcL* and *matK*, which have a greater discriminatory ability than *trnL* (Hollingsworth et al., 2011), may provide increased species-level resolution. However, these regions are much longer and are difficult to recover from degraded samples (Fahner et al., 2016), in addition to being subject to the same limitations of relying on a single gene region. These shortcomings highlight that a more effective approach is needed to fully utilise plant DNA from environmental samples and obtain detailed information on plant community composition.

Targeted capture (also referred to as hybridisation capture) offers an alternative way to overcome the limitations of metabarcoding and bypass the issues associated with PCR amplification (Lemmon & Lemmon, 2013). Targeted capture uses biotinylated RNA molecules called ‘baits’ that are specifically designed to bind to target DNA regions which are then separated from non-target sequences using a magnet These baits eliminate the need for universal primers and, compared to methods such as genome skimming or shotgun sequencing, the targeted nature enhances recovery of organisms of interest within a DNA mixture. This approach reduces overall sequencing costs and increases the amount of information that can be recovered compared to metabarcoding (Foster et al., 2020). Targeted capture approaches have been shown to recover more taxa from environmental samples compared to metabarcoding for a range of different organisms (Dowle et al., 2016; Shokralla et al., 2016; Murchie et al., 2020), and given the potential to recover a large amount of sequence data from multiple gene regions, this approach is also likely to improve taxonomic assignment of sequences from various plant communities.

Here, we employed a targeted capture approach to characterise flowering plant communities in environmental samples, using a bait set designed to simultaneously capture 20 chloroplast regions across all flowering plants (angiosperms). By incorporating multiple gene regions, we aimed to accurately reconstruct plant communities in environmental samples to species level identification, removing the reliance on a single, short barcode. The bait set utilised in this study has not previously been applied to environmental samples and it is rare that studies are undertaken targeting multiple gene regions in a single assay from complex DNA mixtures (one that we know of; Murchie et al., 2020) and never with this many gene regions. We conducted sensitivity and discriminatory power tests on artificial DNA mixtures to assess the threshold of detection and the level of taxonomic resolution offered by the approach before analysing soil samples from a study site where we could verify results using observations from the area.

## 2 Methods

Three trials were conducted to determine the sensitivity, discriminatory power and demonstrate a proof-of-concept for the multi-gene region targeted capture of chloroplast DNA proposed in this study. Each test consisted of a separate experimental setup (see section 2.1) and data analysis (section 2.5) but the same library preparation (section 2.2), multi-gene region bait capture (section 2.3) and read processing and mapping (section 2.4) were conducted for all three trials (Figure 1). Control samples (blanks) were run with each trial sample set to monitor potential contamination and false-positive taxonomic assignments (Ficetola, Taberlet & Coissac, 2016).

**Figure 1.**
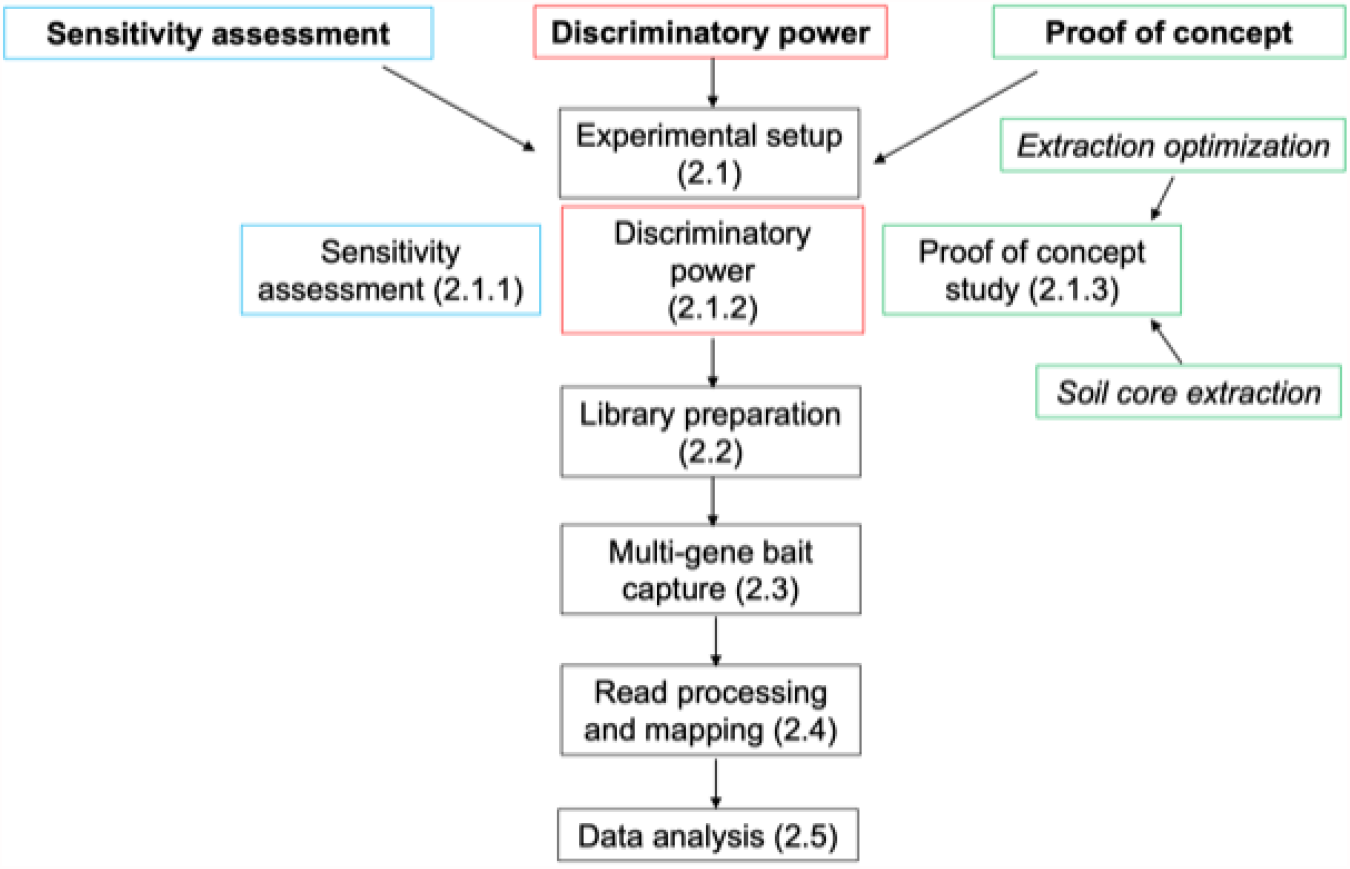
Flow diagram outlining the different tests for this study where colours indicate shared methods.

### 2.1 Experimental setup

#### 2.1.1 Sensitivity assessment

DNA was extracted from three coastal plant species (*Avicenna marina* (grey mangrove), *Tecticornia flabelliformis* (saltmarsh/samphire) and *Zostera marina* (seagrass) using the Plant DNeasy Mini Kit (QIAGEN) as per manufacturer’s instructions and quantified using a Quantus™ Fluorometer and QuantiFluor® dsDNA System (Table S2). An artificial DNA mixture was prepared by combining 40 µL of each DNA extract into a single stock solution (i.e., species were standardised by volume rather than concentration to mimic an environmental sample) which was quantified as above (5.9 ng/µL). This stock was then diluted to the following concentrations: 1 ng/µL, 0.1 ng/µL, 0.01 ng/µL, 0.001 ng/µL and 0.0001 ng/µL. Three replicates of each stock concentration (N=15, volume = 100 µL) were sonicated using a Diagenode Bioruptor® Pico to a size distribution peaking around 400–600 bp (cycle of 15 s on, 90 s off and repeated 7 times to obtain the required size distribution). Further sample processing followed the description in section 2.2 (Figure 1).

#### 2.1.2 Discriminatory power

Total genomic DNA was extracted from 10 coastal plant species (Table S2) using the Plant DNeasy Mini Kit (QIAGEN) following the manufacturer’s instructions. DNA extracts were quantified individually using a Quantus™ Fluorometer and QuantiFluor® dsDNA System. Five different artificial mixtures (with a total volume of 90 µL) were prepared to include an increasing number of species ranging from 3 to 10, standardised by volume rather than concentration (Table 1). The total DNA concentration of each artificial mixture was quantified as above (5.3 ng/µL, 3.85 ng/µL, 3.06 ng/µL, 5 ng/µL, 5.3 ng/µL, respectively) and then standardised to 2 ng/µL. Three replicates from each mixture (N=15, volume = 90 µL) were sonicated using a Diagenode Bioruptor® Pico to a size distribution peaking around 400–600 bp (cycle of 15 s On, 90 s Off and repeat 7 times to obtain the required size distribution). Further sample processing then proceeded to section 2.2 (Figure 1).

**Table 1.**
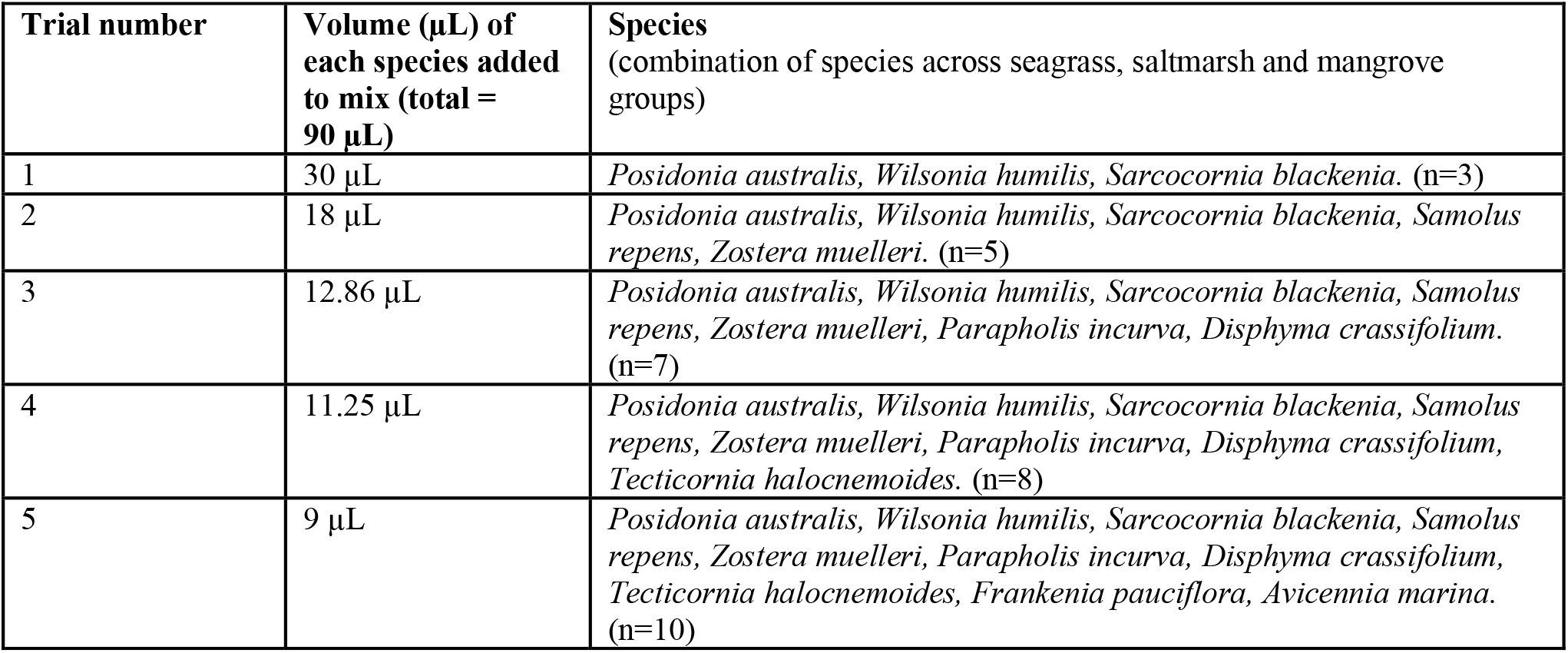
Species mixtures and volumes for DNA artificial mixtures to test discriminatory ability of the targeted capture approach.

### 2.1.3 Proof of concept

#### 2.1.3.1 Optimisation of plant DNA extraction from soils

To optimise the extraction protocol and ensure high plant DNA yield from soil samples, a soil core was collected using a PVC pipe (30 cm long, 75 mm wide) from a shallow landlocked saltwater estuary in Adelaide, South Australia (34° 52.4722’ S, 138° 29.1212’ E). This core was transported vertically to the laboratory and stored at 4°C before sampling in laboratory conditions where surfaces, equipment and solutions were decontaminated prior to use. All laboratory benches were wiped with bleach, ethanol and water and equipment was cleaned with 5% Decon 90 and UV sterilised for 5 min. The sediment core was carefully pushed out of the PVC pipe casing and ∼1 cm of the outer surface was discarded in case of contamination from the PVC pipe. A random location within the top 5 cm of the core was chosen and eight 250 mg samples were collected from the centre of the core for DNA extraction. The total DNA yield of the soil samples was compared between four extraction kits (2 replicates per kit); 1) DNeasy PowerLyzer PowerSoil Kit (QIAGEN^®^), 2) DNeasy Plant Mini Kit, (QIAGEN^®^) 3) QIAamp DNA Stool Mini Kit (QIAGEN^®^) and 4) DNeasy Plant Mini Kit (QIAGEN^®^) combined with the InhibitEx Buffer step from the QIAamp DNA Stool Mini Kit to improve inhibitor removal, hereafter termed ‘combination kit’. The two kits with the highest total DNA yield were then tested specifically for efficiency of plant DNA extraction using plant material homogenised with soil and qPCR (see supplementary methods).

#### 2.1.3.2 DNA extraction from soil core

A single sediment core (75 mm wide and 1 m long) was collected as part of a separate study from an intertidal wetland location on Torrens Island, South Australia (34° 47.574’ S 138° 31.59’ E) in a location where *Avicennia marina* (grey mangrove) was growing (personnel observation). Core transportation, storage and laboratory decontamination was as described above. Two 250 mg samples (A and B) were collected from the centre of the core at 2.5 cm down the length of the core and DNA was extracted using the DNeasy PowerLyzer PowerSoil Kit (QIAGEN^®^) with zirconia beads (based on results from 2.1.3.1). In this instance, sonication was not conducted before library preparation as DNA was assumed to be already fragmented due to the degraded nature of DNA in soils (Corinaldesi, Beolchini & Dell’anno, 2008) and therefore, DNA extracts follow the procedure described in section 2.2 (Figure 1).

### 2.2 Library preparation

An aliquot of the DNA extract was placed into the NEBNext Ultra II Library preparation kit (New England Biolabs^®^) following manufacturer’s instructions with the following modifications: ^1^/3 the recommended reaction volume (16.7 µL) and custom-made stubby (incomplete, P5 and P7 indexes missing) Y-adaptors (25uM) (Glenn et al., 2019) were used at the ligation step. Each adaptor had a unique 8 nucleotide barcode, giving each sample a unique pair of identical internal molecular identifiers (identified as the 8 first base calls for each read). Following adapter ligation, libraries were amplified to detectable concentrations using the supplied Q5 Master Mix at the original reaction volume of 50 µL with in-house primers P7 preCap Long and P5 preCap Long (Cycling conditions: [98°C 10 s, 65°C 30 s, 72°C 30 s] x 17 cycles, 72°C 120 s, 4°C hold). 2 µL of each uniquely indexed library was then visually checked using gel electrophoresis (1 x TE buffer, 1.5% agarose gel for 40 minutes at 80V) and pooled according to concentration estimates (determined via visual inspection) into batches of 8 samples and then purified using AMPure XP (at 0.8 x volume concentration) to remove remaining primers and other impurities. Pooled libraries then progressed to section 2.3.

### 2.3 Multi-gene region bait capture

#### 2.3.1 Bait design

We used the RefSeq release of plastid sequences (https://ftp.ncbi.nlm.nih.gov/refseq/release/plastid/: accessed Oct 2017) to design probes targeting a set of chloroplast gene region regions for angiosperms (Table S1). Using *Arabidopsis lyrata* (Genbank reference NC_034379) as a reference, target regions were extracted from the RefSeq data using Blast (blastn, *e* value < 1e^-50^) and were clustered using CD-HIT (Li & Godzik, 2006) with a 95% identity cut-off, retaining the longest sequence per cluster for probe design. A total of *c*. 2800 representative sequences, ranging in length from 180–900bp (mean 370bp) were used to design *c*. 15,000 120-mer probe sequences with 2X tiling (i.e., each probe overlaps half its length).

### 2.3.2 Targeted capture

Targeted capture was performed on each batch of libraries following the myBaits® Targeted NGS Manual Version 4.01 as per the manufacturer’s instructions. The hybridisation temperature/time was 55°C for 48 hours. Following hybridization, the product was amplified using custom P7 and P5 indexed primers designed in-house using cycling conditions: 98°C 120 s, [98°C 20 s, 60°C 30 s, 72°C 45 s] x 17 cycles, 72°C 30 s, 4°C hold. The final product was an Illumina library where each sample had a unique combination of identical internal dual barcodes (incorporated during library preparation) and two indexes (incorporated after hybridisation). Within our laboratory, all dual barcode-Index 1-Index 2 combinations are only used once, thus reducing contamination risk.

Following targeted capture and amplification, the resulting libraries were run on a 2100 Bioanalyzer (Agilent) using the high sensitivity DNA assay and molarity was calculated between 300–800 bp. All libraries were then pooled in equimolar concentration and purified using AMPure XP (New England Biolabs) at 0.7 x concentration to remove primer dimer and short sequences. The final library, which included all samples in this study, underwent further size selection using a Pippin Prep (Sage Science) with a 1.5% agarose gel cassette set to select between 300–600 bp. The resulting library was quantified using a QuantStudio 6 Flex Real-Time PCR (Thermo Fisher Scientific), diluted to 1.5 nM in 30 µL and sent to the Garvan Institute of Medical Research (Sydney, Australia) to be sequenced on one lane of an Illumina HiSeq X Ten using 2×150 chemistry.

### 2.4 Read processing and mapping

Raw sequences were demultiplexed based on indexes using Illumina Bcl2fastq v2.18.0. The output Read 1 and Read 2 fastq.gz files were then demultiplexed based on the Y-adapter internal barcodes using AdapterRemoval v2 (Schubert, Lindgreen & Orlando, 2016). PALEOMIX (Schubert et al., 2014) was then used to trim adapters (using AdapterRemoval), discard singletons and sequences less than 25 bp, and trim for ambiguous nucleotides and low-quality base calls. BWA-MEM aligner (Li, 2013) was selected within PALEOMIX as the mapping tool while discarding unmapped reads. A specific reference database was used at this step for the different trials, a ‘restricted’ and ‘wider’ reference databases were used in both the sensitivity and discrimination trials and the ‘wider’ reference database was used for the proof of concept (See supplementary information for a description of these databases). Following mapping, Picard Mark Duplicates (Version 4.0.10.1) was used to remove clonality (duplication in read alignment) and the resulting BAM files were then used to generate VCF files using SAMtools mpileup (Li, 2011), specifying ploidy as 1 (as haplotypic organellar DNA) and filtering for base quality < 30, mapping quality < 30 and depth < 50.Variant calls were normalised with BCFtools norm (Li, 2011) and the consensus caller in BCFtools was then used to call final consensus FASTA files, outputting variants with N’s. Output FASTA files were then filtered for length <100bp.

### 2.5 Data analysis

For the sensitivity analysis, the number of chloroplast gene regions recovered in each mixture was counted and calculated as a percentage of the total number of regions that were targeted. We then fitted candidate quasibinomial distributed Generalised Linear Models (GLM) with species, concentration and type of reference as explanatory variables and the percentage of chloroplast gene regions recovered as the response variable. We included different interaction terms between explanatory variables in each candidate GLM and selected the candidate with the greatest model performance (i.e., lowest Akaike Information Criterion). We then conducted an ANOVA on the selected model output to test for significant effects of species, concentration, type of reference and the interactive effect of species and concentration on chloroplast gene region recovery (Table S6).

In the discriminatory power analysis, we found that we could still detect all target gene regions for each of the known species up to the 10 species mixture (the maximum number of species in our tested sample mixtures), indicating that there was no decline in gene region recovery as the number of species in the sample increased from 3 to 10. Therefore, we proceeded to focus our subsequent analyses only on the 10 species sample mixture results. We collated scores of presence or absence for each species, gene region and replicate when sequence reads were mapped to the restricted reference database We then mapped this same 10 species mixture to the wider reference database and combined all replicates to conduct further analyses to observe the level of taxonomic classification achieved for each gene region. We combined replicate FASTA samples (from section 2.4) and used CD-HIT-EST (Li & Godzik, 2006; Fu et al., 2012) to cluster sample sequences with the wider reference database at 95% similarity cut off, removing any samples that did not cluster. We then wrote a custom script in R version 3.5.1 (R Core Team 2018), to determine the lowest discernible taxonomic rank for clusters containing sample sequences (see supplementary methods for details on this). To assess the rate of false-positive assignments we separated these results into 1) species we put into the sample and 2) species we did not put into the sample.

For the proof-of-concept study, we analysed the FASTA files from section 2.4 as above, using the CD-HIT-EST analysis and custom R script.

## 3 Results

### 3.1 Sensitivity assessment

Using artificial DNA mixtures of decreasing concentration, we identified a minimum detection threshold of 0.001– 0.0001 ng/µL total DNA concentration (Figure 2), where below 0.001 ng/µL, the number of target regions recovered across all species was zero. The number of filtered and mapped reads reflects a similar result, where there is a steady decrease in reads with decrease in DNA concentration, and a substantial decline after 0.001 ng/µL (Table S3). There was no difference in target gene region recovery between the wider and restricted reference databases (P=0.85), however, there was an interactive effect between species and concentration (P<0.05) where the percentage of target regions recovered for *Zostera marina* started decreasing at higher concentrations than the other two species (see supplementary Table S6).

**Figure 2.**
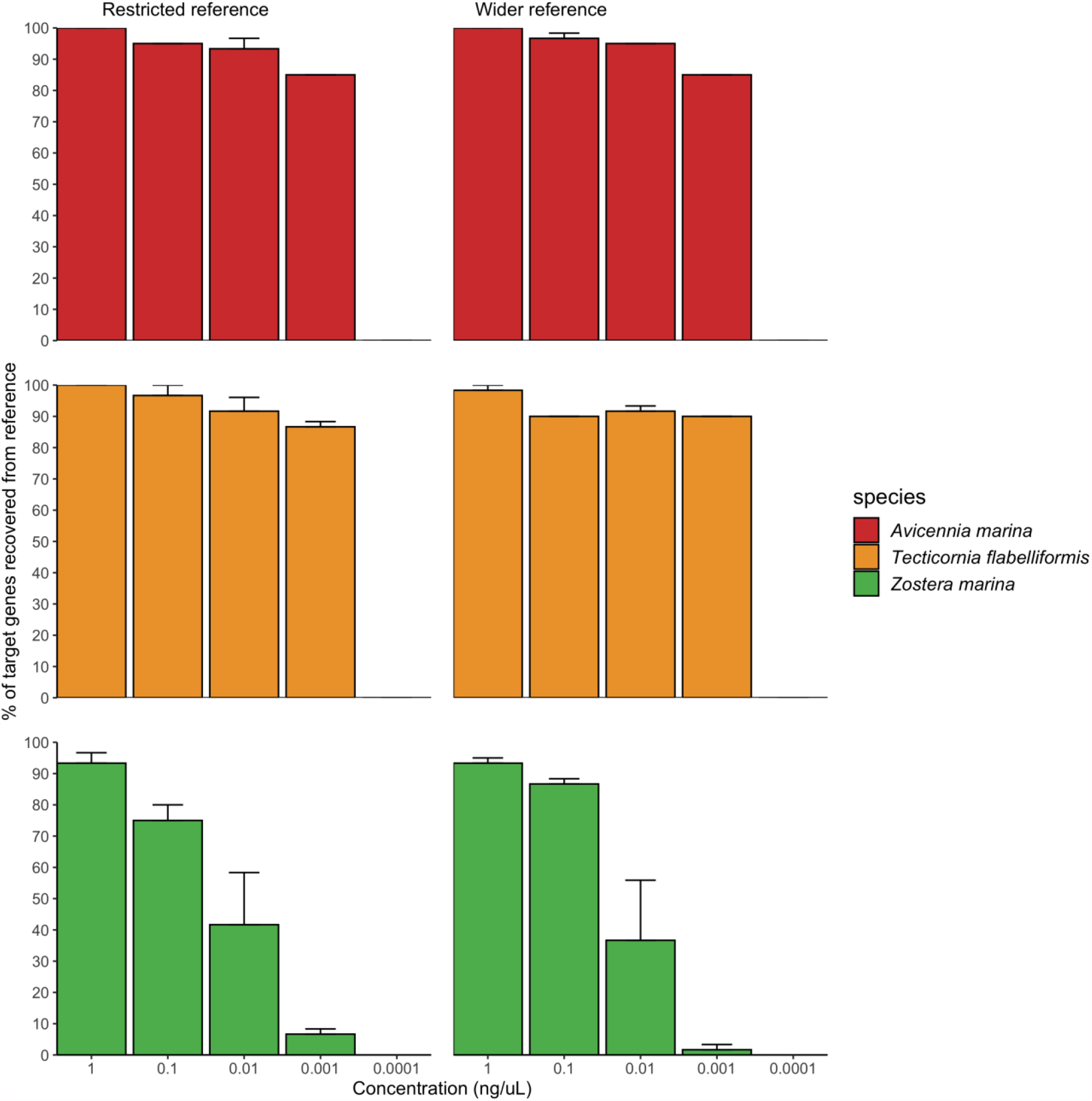
Minimum detection threshold of the targeted capture approach using artificial plant DNA mixtures. All species were in the same mixture but are graphed separately. Concentration reflects the total concentration of the three species mixture, where each species was added in the same volume with a different starting concentration. Restricted and wider reference libraries refer to the different reference databases used in the mapping step.

### 3.2 Discriminatory power

Mapping the maximum 10 species mixture to the restricted reference database, we found 100% recovery for all target gene regions across all species and replicates with one exception, gene region recovery for *Zostera muelleri* was 95% and of these, only 53% were recovered in all three replicates. When the same 10 species artificial DNA mixture was mapped to the wider reference database, and the replicates were pooled, *psbE, atpI* and *rpl16* were recovered across all 10 species. The standard barcoding regions, *matK and rbcL* as well as *atpH, psbD* and *petA*, were recovered across 9 of the 10 species and of these, *matK* had the greatest discriminatory ability, discerning all 9 detected samples to family level or below. Whilst recovered well, the gene region *psbE*, had the lowest discriminatory ability, only capable of discerning samples to order level. Six of the 10 species placed in the mixture were recovered at species level resolution (*Avicennia marina, Disphyma crassifolium, Frankenia pauciflora, Parapholis incurva, Samolus repens and Wilsonia humils)* and the other 4 were recovered at genus level *(Posidonia australis, Sarcocornia quinqueflora, Tecticornia halocnemoides, Zostera muelleri*). In addition to the species that were put in the mixture, additional FASTA files were generated for species that were not placed in the mixture. Taxonomic classification using the clustering algorithm in section 2.5, showed majority of these resolved to order, family or genus of our known species. Figure S2 shows the results of this, where the orders Alismatales, Caryophyllales, Lamiales and Poales are of our known species, as are the families; Aizoaceae, Chenopodiaceae, Poaceae and Primulaceae. Gene regions resolving to the family Scrophulariaceae, the genus Chenopodium and the species’ *Tecticornia flabelliformis, Tecticornia pruinosa and Tecticornia syncarpa* were all recovered but were not placed in the mixture.

### 3.3 Proof of concept

#### 3.3.1 DNA extraction optimisation

The DNeasy PowerLyzer PowerSoil Kit (QIAGEN^®^) showed the highest total DNA yield per mg of sediment compared to the DNeasy Plant Mini Kit, QIAamp DNA Stool Mini Kit and the combination kit (Table S4, P < 0.001). When a known amount of plant material was homogenised with sediment, the DNeasy PowerLyzer PowerSoil Kit (QIAGEN^®^), with zirconia beads replacing the standard glass beads, also yielded significantly more plant DNA than the DNeasy Plant Mini Kit (Table S5; P < 0.05) regardless of plant type (P = 0.053). There was, however, a higher recovery for *Tecticornia flabelliformis* than the other two plant types using the DNeasy PowerLyzer PowerSoil Kit (QIAGEN^®^).

### 3.3.2 Proof of Concept study

Testing the multi-gene region capture approach developed in previous sections on an environmental sample from a coastal wetland, we were able to recover multiple gene regions and species from sediment samples containing an unknown composition of plant genetic material. Figure 4 shows the results of this trial combining the two replicate samples A and B. 11 target gene regions recovered *Avicennia marina* to species level and 9 gene regions resolved to the order Lamiales which is the order *Avicennia marina* belongs to. In addition, two gene regions were recovered for the family Scrophulariaceae. Based on these results, we can conclude that that *Avicennia marina* is present within the environmental samples tested and traces of evidence for the presence of species belonging to the family Scrophulariaceae. Observational data was able to verify this result.

**Figure 3.**
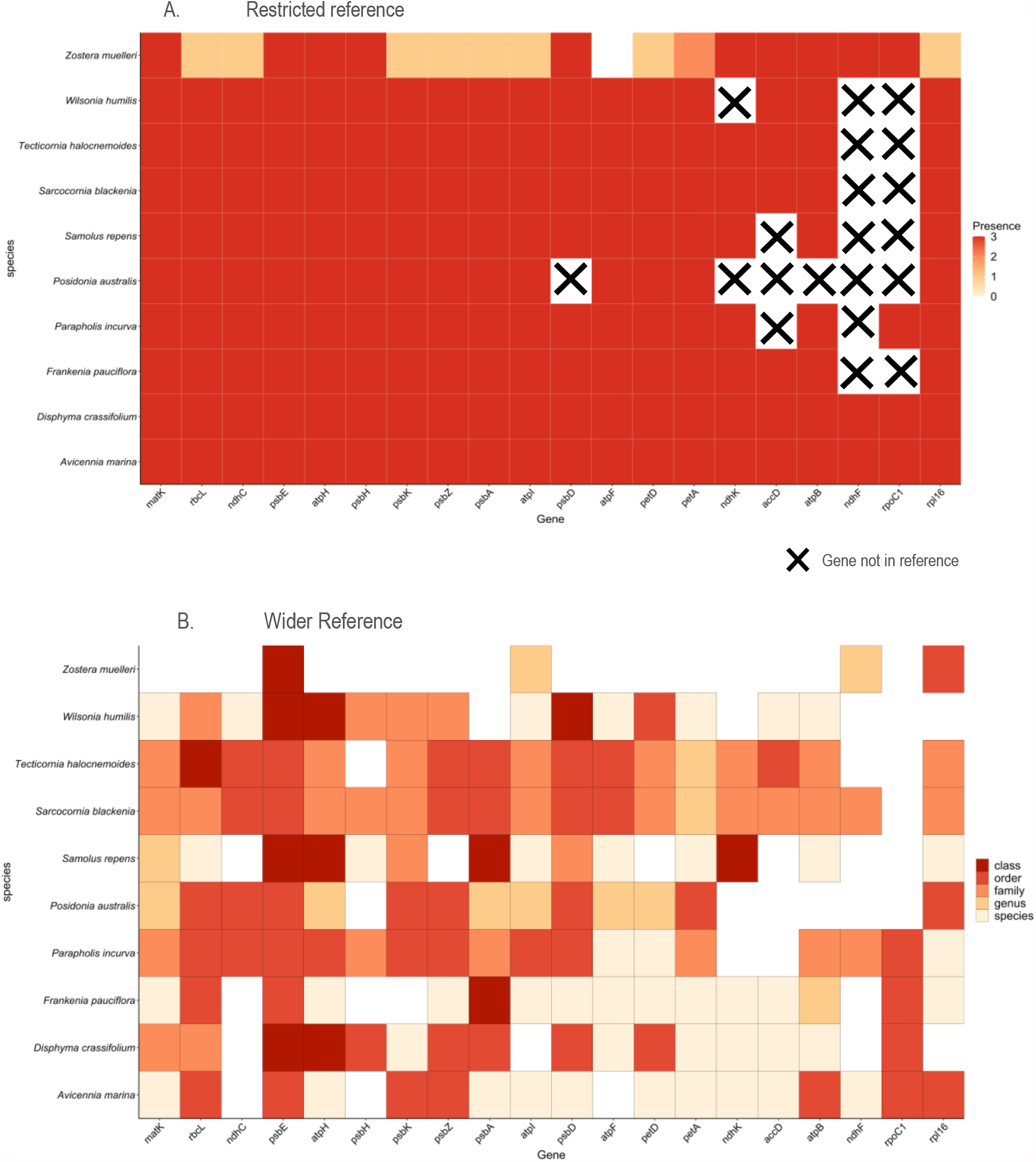
A) The 10 species DNA mixture mapped to a restricted reference library containing only the species included in the mixture. Presence is whether the target gene region was recovered across replicates; B) Taxonomic level of classification when the 10 species mixture is mapped to the wider reference library. All replicate samples are combined.

**Figure 4.**
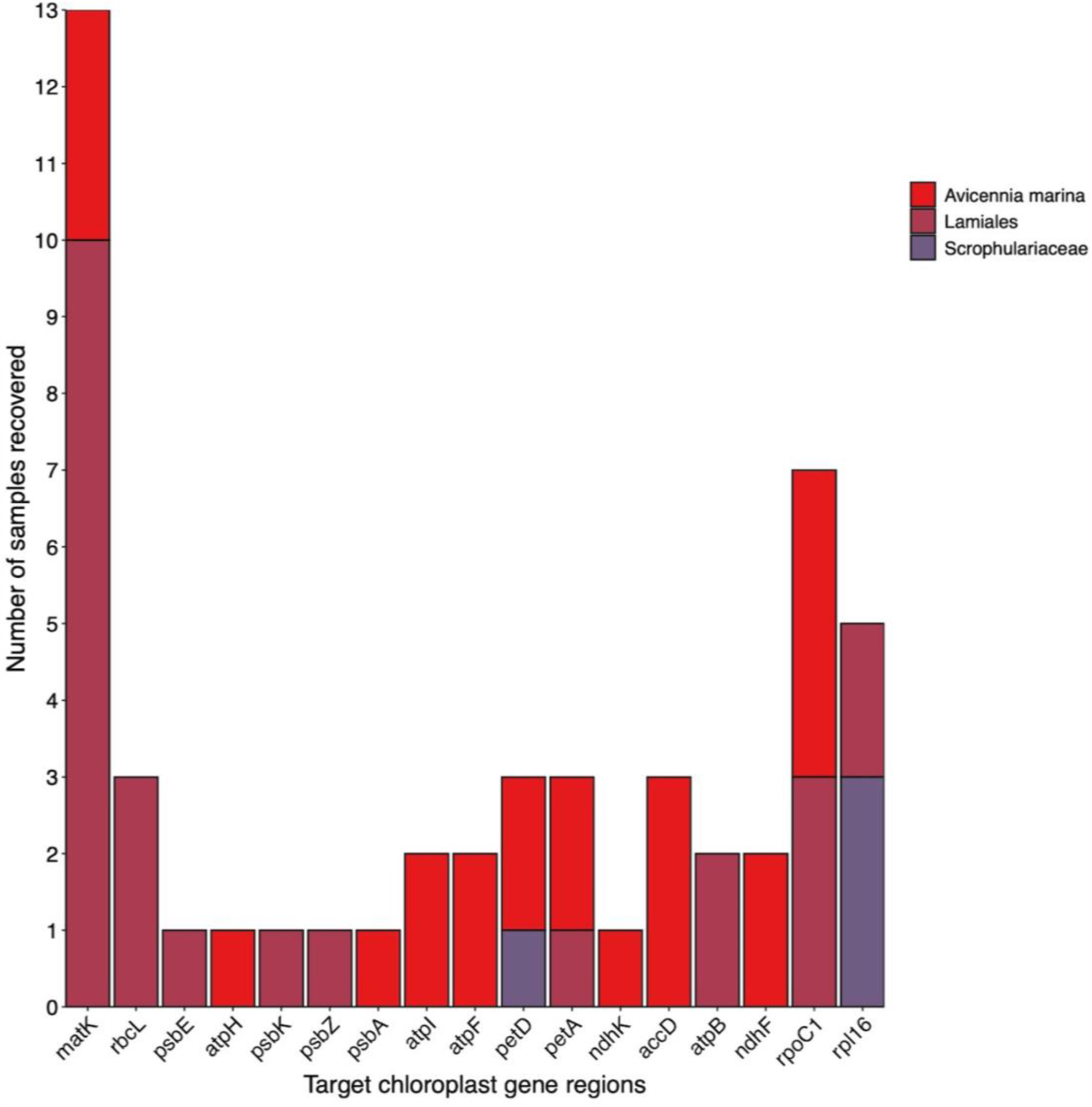
Target chloroplast gene region recovery for species detected in an unknown sediment sample.

### 3.4 Controls

Sample processing and bioinformatic analysis of sample blank controls for each stage of this study yielded no matches to either the restricted or wider reference databases and therefore we are confident there has not been any contamination in this study.

## 4 Discussion

This study presents a novel approach to detecting and characterising flowering plant DNA in environmental samples. We have demonstrated this targeted capture technique can recover multiple chloroplast gene regions from environmental samples in a single assay. We show that the lowest detection level of this approach is down to 0.001 ng/µL and therefore, this method is sensitive enough to detect low concentrations of DNA that are likely to be found in environmental samples due to the degradation that occurs (Pedersen et al., 2015). Furthermore, we highlight that multi-gene recovery is possible across many taxa (i.e., up to 10 species in the same sample), however, the comprehensiveness of reference databases used in the read mapping step can influence the number of gene regions recovered. Here, we also showed that the target chloroplast gene regions have different levels of discrimination for different flowering plant groups and are hence a multi-gene approach increases the likelihood of recovering all species present in a sample at either species or genus level. Applying this method to in-situ soil samples showed that optimal plant DNA extraction from soils can be achieved using the DNeasy PowerLyzer PowerSoil Kit (QIAGEN^®^), which aligns with other studies that confirm the utility of this kit for maximum yield and greatest biodiversity detection amongst available methods (Hermans, Buckley & Lear, 2018). We also successfully identified the mangrove species, *Avicennia marina* as being present in this sample, based on the recovery of multiple target gene regions that resolved to this species.

### 4.1 Detection threshold when targeting multiple regions and species in environmental samples

Using the multi-gene region targeted approach, our ability to detect all species present in an artificial mixture was significantly impacted by total DNA concentration of the sample. In our sensitivity analysis, the percentage of target gene regions recovered for *Zostera marina* slowly declined below 1 ng/µL to no recovery at 0.0001 ng/ µL, whereas the other two species (*Tecticornia flabelliformis* and *Avicennia marina*) recorded a decline only below 0.001 ng/µL. We attribute this result to the varying DNA input concentrations of species in this trial where *Z. marina* was present at lower concentration in the starting mixture, contributing to only 1% whereas *A. marina* and *T. flabelliformis* contributed 46% and 53% respectively (Table S2). However, despite the lower initial concentration, we were still able to recover >10% of *Z. marina* target chloroplast regions down to 0.001 ng/µL. The sharp decrease in gene region recovery for all species in the mixture after 0.001 ng/µL implies that total DNA concentration significantly impacts gene region recovery for all species in a mixture, regardless of the amount of DNA contributed for each individual species. As environmental DNA and ancient samples have characteristically degraded DNA and thus low concentration (Pedersen et al., 2015), a multi-gene region targeted approach seems the best option to increase the chances of capturing the entire plant community present.

### 4.2 The power of a multi-gene region approach

We have demonstrated that the multi-gene region capture method developed in this study can successfully recover multiple regions across many species when we have prior knowledge of what is in a sample (Figure 3A). The lower gene region recovery for *Zostera muelleri* in Figure 3, can again be attributed to the low input DNA concentration (Table S2). However, the fact that both Zostera species in each of our trials had the lowest number of recovered regions, suggests additional factors may be influencing the amount of DNA recovered for this genus, such as the chloroplast copy number (Sakamoto & Takami, 2018). This highlights the possibility that taxa within environmental samples can have naturally unequal amounts of DNA which can potentially skew the number and type of gene regions recovered for different plant groups. Therefore, targeting a single region, such as in standard metabarcoding, may give an inaccurate picture of the plant community composition present in a sample as some species will be missed.

Furthermore, when we then mapped the same 10 species mixture to the wider reference library, we found that, for some gene regions, reads were no longer mapped directly to the expected taxon. It is likely, for these regions, mapping has occurred to related species with a reference sequence for that species that it couldn’t discriminate between. This highlights how important a comprehensive reference database is when trying to decipher plant community presence in environmental samples. Having confidence that the read mapping step has correctly assigned reads to the reference database of species’ that are present in the sample is important. Furthermore, as we showed in Figure 3B, the variability and thus discrimination potential varies between gene regions and between flowering plant groups. This is why we undertook additional analyses to assign taxonomic classification to sample sequences, so we could ensure the sequence was unique to that species and to increase confidence when assigning species’ to unknown sequences. We assigned species presence for 6 of the 10 species we placed in the mixture and the rest were assigned to genus level.

The additional species detected that were not placed in the mixture is concerning and highlights possible shortcomings of our target bait set. Detecting *Tecticornia flabelliformis, Tecticornia pruinosa and Tecticornia syncarpa*, despite only putting *Tecticornia halocnemoides* in the mixture is likely due to the poor discrimination of these taxa for this genus. There are several explanations for the detection of other *Tecticornia* species in these samples. First, is that the species boundaries are poorly resolved with chloroplast-based data sets (e.g., Shepherd et al., 2004).

Alternatively, these taxa may be forming hybrids, an observation consistent with other data sets under study (E. Biffin and M. Waycott pers. comm.). For either hypothesis, additional chloroplast-based gene regions or the addition of nuclear gene regions may help to separate this difficult genus. Unfortunately, due to these factors, we cannot tease apart the exact cause of these false positive detections but as they are of the same genus as the species we placed in the mixture, it is unlikely there is something wrong with the method. This is similar to the detection of the family Scrophulariaceae and the genus *Chenopodium*, while these were not the samples placed in the mixture, they are members of the same family and genus and may be assigned due to an inability of the markers to discriminate at the species level.

Using the 20 chloroplast gene regions targeted in this study, we were able to accurately determine the presence of *Avicennia marina* in an unknown environmental sample, and detect the family Scrophulariaceae and the order Lamiales. The subsequent verification of the presence of *Avicennia marina* through observation, demonstrates the power in multiple gene regions to accurately detect flowering plant species within environmental samples. We acknowledge that metabarcoding can be conducted for many gene regions in order to increase the genetic information required to confidently assign taxonomy, however, this cannot be done in a single assay nor in a targeted way to increase efficiency. Therefore, the targeted approach demonstrated here, is more prudent to use when recovering genetic information from environmental samples across many plant groups, where this can even be implemented beyond simple species identification but to other fields such as phylogenetic reconstruction (Adams et al., 2019).

### 4.3 Future directions

The the limitations of the universal barcode for plants means capturing the genetic variation required to disentangle species concepts across all flowering plant groups is difficult with a single, or even a few, gene region. Hence, we have turned to targeting multiple regions in a single assay to create redundancy in our detection ability as we do not rely on one region but 20, to provide evidence of species presence. Future studies can be done in this area to improve bioinformatic analyses to place certainty parameters around species presence based on the number of target gene regions recovered and the level of discrimination they offer i.e., their information content. In addition, developing a bait set to target additional chloroplast and/or nuclear gene regions, will improve species detection. Additionally, modifying either the hybridisation time or lowering the temperature, could help recover more DNA from environmental samples. This may improve the amount of data generated for each gene region, or recover more regions, which would increase the amount of data generated for each species. This will improve the ability to disentangle species concepts in unknown mixtures and form more accurate conclusions around flowering plant presence in environmental samples. This can then be used to identify whether species have been lost from a system or detect invasive species as well as track food webs and reconstruct past flowering plant communities.

### 5 Conclusion

The proliferation of environmental DNA studies highlights a growing interest in methods for environmental monitoring (Beng & Corlett, 2020). While current DNA metabarcoding using a single gene region provides a tool for monitoring our natural environments (Balint et al., 2018), the application and interpretation of this data relies on accurate taxonomic identification of sequences recovered from an environmental sample (Pedersen et al., 2015). We have shown that a targeted capture approach has the ability to improve both the recovery of multiple species in a mixture and assign taxonomy across a large number of flowering plant groups. Obtaining a suite of genetic data across diverse plant taxa, equips us with the ability to generate reliable conclusions regarding plant communities in environmental samples, which can be used to improve monitoring, management and conservation outcomes. This study is the first to apply a targeted capture approach to recover multiple gene regions from environmental samples in a single assay, and thus enables a more reliable and accurate method to determine plant presence in environmental samples.

## 6 Authorship statement

The authors declare that the research was conducted in the absence of any commercial or financial relationships that could be construed as a potential conflict of interest.

## 7 Author Contributions

NF, JY and MW conceived the ideas. NF, JY, MW, KV and EB designed the methodology. NF collected the data and ran the experiments. NF and VT analysed the data, MW advised on interpretation of analysis and results. NF wrote the manuscript. All authors contributed to editing and preparing the manuscript.

## 9 Acknowledgements

This research was undertaken on the lands of the Kaurna people. We acknowledge and respect their spiritual relationship with their country, their cultural and heritage beliefs and their rights as the Traditional Owners of the land. We would like to thank Lucy and Emma for their assistance with field work.

## 10 Data Availability Statement

All raw, demultiplexed files are available on Figshare with the DOI: 10.25909/15049151. Sediment samples are available on Sequence Read Archive server with the BioProject ID: PRJNA749388

## References

Adams, C. I. M., Knapp, M., Gemmell, N. J., Jeunen, G. J., Bunce, M., Lamare, M. D., et al. (2019). Beyond Biodiversity: Can Environmental DNA (eDNA) Cut It as a Population Genetics Tool? Regions (Basel), 10. 192

Balint, M., Pfenninger, M., Grossart, H. P., Taberlet, P., Vellend, M., Leibold, M. A., et al. (2018). Environmental DNA Time Series in Ecology. Trends in Ecology & Evolution, 33, 945–957.

Beng, K. C. & Corlett, R. T. (2020). Applications of environmental DNA (eDNA) in ecology and conservation: opportunities, challenges and prospects. Biodiversity and Conservation, 29, 2089–2121.

Brown, S. K. & Blois, J. L. (2001). Ecological Insights from Ancient DNA. English Literary Studies Monograph Series, 1–7.

Corinaldesi, C., Beolchini, F. & Dell’anno, A. (2008). Damage and degradation rates of extracellular DNA in marine sediments: implications for the preservation of gene sequences. Molecular Ecology, 17, 3939–51.

Del Carmen Gomez Cabrera, M., Young, J. M., Roff, G., Staples, T., Ortiz, J. C., Pandolfi, J. M., et al. (2019). Broadening the taxonomic scope of coral reef palaeoecological studies using ancient DNA. Molecular Ecology, 28, 2636–2652.

Dowle, E. J., Pochon, X., J, C. B., Shearer, K. & Wood, S. A. (2016). Targeted gene enrichment and high-throughput sequencing for environmental biomonitoring: a case study using freshwater macroinvertebrates. Molecular Ecololgy Resources, 16, 1240–54.

Epp, L. S., Boessenkool, S., Bellemain, E. P., Haile, J., Esposito, A., Riaz, T., et al. (2012). New environmental metabarcodes for analysing soil DNA: potential for studying past and present ecosystems. Molecular Ecology, 21, 1821–33.

Fahner, N. A., Shokralla, S., Baird, D. J. & Hajibabaei, M. (2016). Large-scale monitoring of plants through environmental DNA metabarcoding of soil: recovery, resolution, and annotation of four DNA markers. PLoS ONE, 11, e0157505.

Ficetola, G. F., Taberlet, P. & Coissac, E. (2016). How to limit false positives in environmental DNA and metabarcoding? Molecular Ecololgy Resources, 16, 604–7.

Fordham, D. A., Akçakaya, H. R., Alroy, J., Saltré, F., Wigley, T. M. L. & Brook, B. W. (2016). Predicting and mitigating future biodiversity loss using long-term ecological proxies. Nature Climate Change, 6, 909–916.

Foster, N. R., Gillanders, B. M., Jones, A. R., Young, J. M. & Waycott, M. (2020). A muddy time capsule: using sediment environmental DNA for the long-term monitoring of coastal vegetated ecosystems. Marine and Freshwater Research, 71.

Fu, L., Niu, B., Zhu, Z., Wu, S. & Li, W. (2012). CD-HIT: accelerated for clustering the next-generation sequencing data. Bioinformatics, 28, 3150–2.

Furlan, E. M., Davis, J. & Duncan, R. P. (2020). Identifying error and accurately interpreting eDNA metabarcoding results: A case study to detect vertebrates at arid zone waterholes. Molecular Ecololgy Resources.20(5), 1259–1276

Giguet-Covex, C., Pansu, J., Arnaud, F., Rey, P. J., Griggo, C., Gielly, L., et al. (2014). Long livestock farming history and human landscape shaping revealed by lake sediment DNA. Nature Communications, 5, 3211.

Glenn, T.C., Nilsen, R.A., Kieran, T.J., Finger, J.W., Pierson, T.W., Bentley, K.E., Hoffberg, S.L., Louha, S., García-De León, F.J., del Rio Portilla, M.A. and Reed, K.D., 2016. Adapterama I: Universal stubs and primers for thousands of dual-indexed Illumina libraries (iTru & iNext). BioRxiv, 049114.

Guindon, S., Dufayard, J.-F., Lefort, V., Anisimova, M., Hordijk, W. & Gascuel, O. (2010). New algorithms and methods to estimate maximum-likelihood phylogenies: assessing the performance of PhyML 3.0. Systematic Biology, 59, 307–321.

Hermans, S. M., Buckley, H. L. & Lear, G. (2018). Optimal extraction methods for the simultaneous analysis of DNA from diverse organisms and sample types. Molecular Ecololgy Resources, 18, 557–569.

Hollingsworth, P. M., Graham, S. W. & Little, D. P. (2011). Choosing and using a plant DNA barcode. PLoS ONE, 6, e19254.

Hollingsworth, P. M., Li, D.-Z., Van Der Bank, M. & Twyford, A. D. (2016). Telling plant species apart with DNA: from barcodes to genomes. Philosophical Transactions of the Royal Society B: Biological Sciences, 371.

Katoh, K., Misawa, K., Kuma, K. I. & Miyata, T. (2002). MAFFT: a novel method for rapid multiple sequence alignment based on fast Fourier transform. Nucleic Acids Research, 30, 3059–3066.

Lemmon, E. M. & Lemmon, A. R. (2013). High-throughput genomic data in systematics and phylogenetics. Annual Review of Ecology, Evolution, and Systematics, 44, 99–121.

Li, H. (2011). A statistical framework for SNP calling, mutation discovery, association mapping and population genetical parameter estimation from sequencing data. Bioinformatics, 27, 2987–93.

Li, H. (2013). Aligning sequence reads, clone sequences and assembly contigs with BWA-MEM. arXiv preprint arXiv:1303.3997.

Li, W. & Godzik, A. (2006). Cd-hit: a fast program for clustering and comparing large sets of protein or nucleotide sequences. Bioinformatics, 22, 1658–9.

Murchie, T. J., Kuch, M., Duggan, A. T., Ledger, M. L., Roche, K., Klunk, J., et al. (2020). Optimizing extraction and targeted capture of ancient environmental DNA for reconstructing past environments using the PalaeoChip Arctic-1.0 bait-set. Quaternary Research, 1–24.

NCBI Resource Coordinators, 2018. Database resources of the National Center for Biotechnology Information. Nucleic Acids Research, 46, D8–D13.

Pansu, J., Giguet-Covex, C., Ficetola, G. F., Gielly, L., Boyer, F., Zinger, L., et al. (2015). Reconstructing long-term human impacts on plant communities: an ecological approach based on lake sediment DNA. Molecular Ecology, 24, 1485–98.

Paradis, E., Claude, J. & Strimmer, K. (2004). APE: Analyses of Phylogenetics and Evolution in R language. Bioinformatics, 20, 289–90.

Parducci, L., Matetovici, I., Fontana, S. L., Bennett, K. D., Suyama, Y., Haile, J., et al. (2013). Molecular-and pollen-based vegetation analysis in lake sediments from central Scandinavia. Molecular Ecology, 22, 3511–24.

Pedersen, M. W., Overballe-Petersen, S., Ermini, L., Der Sarkissian, C., Haile, J., Hellstrom, M., et al. (2015). Ancient and modern environmental DNA. Philosophical Transactions of the Royal Society B: Biological Sciences, 370, 20130383.

R Core Team (2018). R: A language and environment for statistical computing. R Foundation for Statistical Computing, Vienna, Austria. URL https://www.R-project.org/

Reef, R., Atwood, T. B., Samper-Villarreal, J., Adame, M. F., Sampayo, E. M. & Lovelock, C. E. (2017). Using eDNA to determine the source of organic carbon in seagrass meadows. Limnology and Oceanography, 62, 1254–1265.

Ruppert, K. M., Kline, R. J. & Rahman, M. S. (2019). Past, present, and future perspectives of environmental DNA (eDNA) metabarcoding: A systematic review in methods, monitoring, and applications of global eDNA. Global Ecology and Conservation, 17.

Russell V. Lenth (2016). Least-Squares Means: The R Package lsmeans. Journal of Statistical Software, 69(1), 1–33. doi:10.18637/jss.v069.i0

Sakamoto, W. & Takami, T. (2018). Chloroplast DNA Dynamics: Copy Number, Quality Control and Degradation. Plant and Cell Physiology, 59, 1120–1127.

Schubert, M., Ermini, L., Der Sarkissian, C., Jonsson, H., Ginolhac, A., Schaefer, R., et al. (2014). Characterization of ancient and modern genomes by SNP detection and phylogenomic and metagenomic analysis using PALEOMIX. Nature Protocols, 9, 1056–82.

Schubert, M., Lindgreen, S. & Orlando, L. (2016). AdapterRemoval v2: rapid adapter trimming, identification, and read merging. BMC Research Notes, 9, 88.

Shepherd KA, Waycott M, Calladine A (2004). Radiation of the Australian Salicornioideae (Chenopodiaceae): based on evidence from nuclear and chloroplast DNA sequences. American Journal of Botany 91, 1387–1397

Shokralla, S., Gibson, J. F., King, I., Baird, D. J., Janzen, D. H., Hallwachs, W., et al. (2016). Environmental DNA barcode sequence capture: targeted, PCR-free sequence capture for biodiversity analysis from bulk environmental samples. BioRxiv, 087437.

Taberlet, P., Coissac, E., Pompanon, F., Gielly, L., Miquel, C., Valentini, A., et al. (2006). Power and limitations of the chloroplast trnL (UAA) intron for plant DNA barcoding. Nucleic Acids Research, 35, e14–e14.

Thomsen, P. F. & Willerslev, E. (2015). Environmental DNA – An emerging tool in conservation for monitoring past and present biodiversity. Biological Conservation, 183, 4–18.

Willerslev, E., Davison, J., Moora, M., Zobel, M., Coissac, E., Edwards, M. E., et al. (2014). Fifty thousand years of Arctic vegetation and megafaunal diet. Nature, 506, 47–51.

Willerslev, E., Hansen, A. J., Binladen, J., Brand, T. B., Gilbert, M. T. P., Shapiro, B., et al. (2003). Diverse plant and animal genetic records from Holocene and Pleistocene sediments. Science, 300, 791–795.

Yoccoz, N. G., Brathen, K. A., Gielly, L., Haile, J., Edwards, M. E., Goslar, T., et al. (2012). DNA from soil mirrors plant taxonomic and growth form diversity. Molecular Ecology, 21, 3647–55.

Zimmermann, H. H., Raschke, E., Epp, L. S., Stoof-Leichsenring, K. R., Schwamborn, G., Schirrmeister, L., et al. (2017). Sedimentary ancient DNA and pollen reveal the composition of plant organic matter in Late Quaternary permafrost sediments of the Buor Khaya Peninsula (north-eastern Siberia). Biogeosciences, 14, 575–596.

Zinger, L., Bonin, A., Alsos, I. G., Balint, M., Bik, H., Boyer, F., et al. (2019). DNA metabarcoding-Need for robust experimental designs to draw sound ecological conclusions. Molecular Ecology, 1857–1862.

