## Supplementary material for "A multi-gene region targeted capture approach to detect plant DNA in environmental samples: A case study from coastal environments"

### 1 Supplementary methods

#### 1.1 Optimisation of plant DNA extraction from soils

**Total DNA yield*:*** Four DNA extraction methods (two replicates for each) were tested to compare total DNA yield: 1) DNeasy PowerLyzer PowerSoil Kit (QIAGEN®), 2) DNeasy Plant Mini Kit, (QIAGEN®) 3) QIAamp DNA Stool Mini Kit (QIAGEN®) and 4) DNeasy Plant Mini Kit (QIAGEN®) combined with the InhibitEx Buffer step from the QIAamp DNA Stool Mini Kit to improve inhibitor removal, hereafter termed 'combination kit'. For the initial homogenisation step, glass beads were used in the DNeasy PowerLyzer PowerSoil Kit and the QIAamp DNA Stool Mini Kit whereas zirconium beads (1 mm and 2.3 mm) were used in the DNeasy Plant Mini Kit and the combination kit, following kit instructions. DNA yield was determined via Quantus™ Fluorometer (Promega®) and QuantiFluor® dsDNA System in ng/µL then converted to ng/mg of soil. A one-way ANOVA was fitted to these values in R version 3.5.1 (R Core Team 2018) with extraction kit as an explanatory variable and DNA yield as the response variable. Planned contrasts between kits were conducted using the lsmeans package (Lenth & Lenth, 2018) and the Tukey adjustment method. DNA concentration values were square root transformed due to the skewed nature of the results and to satisfy the requirement of normality of residuals.

**Recovery of plant DNA from sediment*:*** To specifically determine the efficiency of plant DNA extraction, soil and plant material were homogenised to observe whether the extraction kit performance improved or changed when there was known plant material in the mixture, and whether plant type influenced DNA recovery. Three plant/soil mixtures were made consisting of ~200 mg of soil (same core and process as above) and ~20 mg of one of the following plant types; *Zostera muelleri*, *Avicenna marina* or *Tecticornia flabelliformis.* Three replicates of each mixture underwent DNA extraction (in triplicate) using only the DNeasy PowerLyzer PowerSoil Kit (QIAGEN®) and the DNeasy Plant Mini Kit, (QIAGEN®) based on the results of the previous trials. Zirconia beads were used in both kits as glass beads could not fully pulverise plant material (personal observation). Total DNA yield was determined via a Quantus™ Fluorometer and QuantiFluor® dsDNA System in ng/µL then converted to ng/mg of plant material. A Shapiro-Wilks normality test was performed, and this satisfied the requirements for normality ($P > 0.05$). A two-way ANOVA was then fitted to the data in R version 3.5.1 (R Core Team 2018) with kit and plant type as explanatory variables and DNA concentration as the response variable.

#### 1.2 Different reference libraries used in read processing and mapping

For the sensitivity and discrimination trials, reads were mapped to both a 'restricted' and a 'wider' reference database. The wider reference database consisted of 94 coastal temperate plant species (Foster *et al.* 2021, unpublished data) generated from voucher specimens using the same chloroplast bait set as applied to the test samples (Table S1) and combined with references from the National Centre for Biotechnology Information (NCBI Resource Coordinators, 2018) database consisting of all available sequences for the 20 target chloroplast genes of South Australian flora.
The restricted database consisted of a subset of this database to only include plant species present in the artificial mixtures in order to quantify the success of the approach in regard to gene recovery.

#### 1.3 Data analysis

**Discriminatory power CD-HIT-EST and custom R script**

FASTA files for both the 10 species artificial mixture and proof-of-concept (section 2.4), were clustered with the wider reference database using CD-HIT-EST (Li & Godzik, 2006; Fu et al., 2012). A flow diagram for this analysis is outlined in Figure S1 including the read processing and mapping steps.

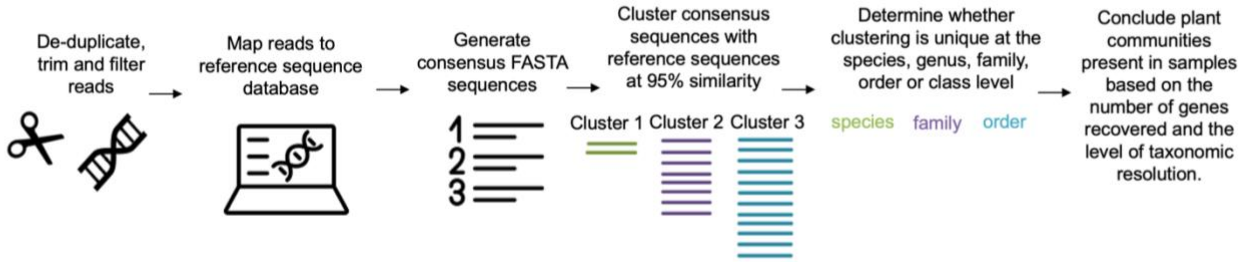


**Figure S1. Flow diagram depicting the series of analytical steps used to determine plant community composition in the discriminatory power trial and the proof-of-concept trial.**

The threshold for sequence identity was set to 0.95, length and clustering of sequences was specified to cluster at the most similar cluster (-g 1) and alignment was set to cover at least 10% of the representative sequence and 90% of the shorter sequence (-aL 0.1 -aS 0.9). A custom script was written in R version 3.5.1 (R Core Team 2018), using packages taxize (Chamberlain and Szocs, 2013), TAI (Drost et al., 2018), dplyr (Wickham et al., 2020), stringi (Gagolewski et al. 2020), stringr (Wickham et al., 2019) and tidyr (Wickham and Henry, 2019). This script unpacked the output.clstr file from CD-HIT-EST and generated upstream taxonomic assignment for each sample, in each cluster. The data was then separated into each cluster and the script identified the highest common taxonomic ranking as an output. Each of the sample sequences were then separated so that the final dataset contained assigned sample taxonomy generated from the mapping of reads (section 2.4) and a ranking for the level of taxonomic clustering this sequence provided. This allowed us to determine whether the read mapping step and subsequent generation of FASTA files from our sample reads was an accurate determination of what was present in the mixture. If the sample sequence clustered broadly with other sequences, and thus the highest common taxonomic level was order or family, it meant either, the gene in question did not have enough discriminatory power/could not discern between species on its own, or the sample sequence reads did not contain enough genetic data to generate informative FASTA sequences i.e. read depth did not meet the assigned threshold to call a base and instead missing data values were inserted (N's) and thus the length of the sequence was too short to be informative. This process helped us eliminate any mapping error and take into account the different discriminatory ability of chloroplast gene regions for the different flowering plant groups.

### 1.4 References


Chamberlain S. & Szocs E.(2013). taxize - taxonomic search and retrieval in R. F1000Research, 2:191. \https://f1000research.com/articles/2-191/v2

Drost HG, Gabel A, Jiu J, Quint M, Grosse I (2018). myTAI: evolutionary transcriptomics with R. *Bioinformatics* doi:10.1093

Fu, L., Niu, B., Zhu, Z., Wu, S. & Li, W. (2012). CD-HIT: accelerated for clustering the next-generation sequencing data. *Bioinformatics,* 28**,** 3150-2.

Gagolewski M. (2020). R package stringi: Character string processing facilities. http://www.gagolewski.com/software/stringi/.

Lenth, R. & Lenth, M. R. (2018). Package ‘lsmeans’. *The American Statistician,* 34**,** 216-221.

Li, W. & Godzik, A. (2006). Cd-hit: a fast program for clustering and comparing large sets of protein or nucleotide sequences. *Bioinformatics,* 22**,** 1658-9.

R Core Team (2018). R: A language and environment for statistical computing. R Foundation for Statistical Computing, Vienna, Austria. URL https://www.R-project.org

Wickham, H., François, R., Lionel H., Müller, K. (2020). dplyr: A Grammar of Data Manipulation. R package version 1.0.2. https://CRAN.R-project.org/package=dplyr

Wickham, H. (2019). stringr: Simple, Consistent Wrappers for Common String Operations. R package version 1.4.0. https://CRAN.R-project.org/package=stringr

Wickham, H., & Henry, L. (2019). tidyr: Tidy Messy Data. R package version 1.0.0. https://CRAN.R-project.org/package=tidyr

### 2 Supplementary Figures and tables

**Table S1. List of target chloroplast gene regions screened for as 'on target' in the bait set used to target capture eDNA.**

| Gene | Name |
|---|---|
| psbA | photosystem II protein D1 |
| matK | maturase K |
| psbK | photosystem II protein K |
| atpF | ATP synthase CF0 subunit I |
| atpH | ATP synthase CF0 subunit III |
| atpI | ATP synthase CF0 subunit IV |
| psbD | photosystem II protein D2 |
| psbZ | photosystem II protein Z |
| rbcL | ribulose-1,5-bisphosphate carboxylase/oxygenase large subunit |
| atpB | ATP synthase CF1 beta subunit |
| accD | acetyl-CoA carboxylase carboxyltransferase beta subunit |
| petA | cytochrome f |
| psbE-psbF | photosystem II cytochrome b559 alpha subunit/photosystem II cytochrome b559 beta subunit |
| psbN-psbH | photosystem II protein N/photosystem II phosphoprotein |
| petD | cytochrome b6/f complex subunit IV |
| rpl14-rpl16 | ribosomal protein L14/ribosomal protein L16 |
| rpoC1 | β subunit of RNA polymerase |
| ndhK | NADH-plastoquinone oxidoreductase subunit K |
| ndhF | NADH-plastoquinone oxidoreductase subunit 5 |
| ndhC | NADH-plastoquinone oxidoreductase subunit 3 |

**Table S2. Summary of DNA concentration (ng/µL) and collection information for samples used in the sensitivity assessment (section 2.1.1 from main text) and discriminatory power (section 2.1.2 from main text)**

| Species | Concentration (ng/µL) | Collection date | Country | Location | Latitude | Longitude | Specimen voucher number |
|---|---|---|---|---|---|---|---|
| *Avicennia marina* | 9.72 | 00/12/2017 | Australia | South Australia St. Kilda | -34.73 | 138.52 | AD284320 |
| *Tecticornia flabelliformis* | 11.35 | 00/02/2018 | Australia | South Australia Middle Beach | -32.87 | 134.11 | AD284322 |
| *Zostera marina* | 0.22 | 14/04/1975 | Australia | Washington, Stanley, Park B.C. | 47.54 | -122.84 | AD283375 |
| *Posidonia australis* | 1.86 | 25/11/2009 | Australia | Rottnest Island, Stark Bay, WA | -32.26 | 115.70 | AD272341 |
| *Wilsonia humilis* | 1.89 | 00/02/2018 | Australia | South Australia Port Gawler | -34.65 | 138.45 | AD284323 |
| *Sarcocornia blackenia* | 4.97 | 00/02/2018 | Australia | South Australia Port Gawler | -34.73 | 138.52 | AD284325 |
| *Samolus repens* | 2.20 | 00/02/2018 | Australia | South Australia Port Gawler | -34.65 | 138.45 | AD284324 |
| *Zostera muelleri* | 0.28 | 20/01/2015 | Australia | Stansbury SA | -34.91 | 137.81 | AD272441 |
| *Parapholis incurva* | 0.57 | 00/12/2017 | Australia | South Australia St. Kilda | -34.73 | 138.52 | AD284314 |
| *Disphyma crassifolium* | 0.63 | 00/02/2018 | Australia | South Australia St. Kilda | -34.73 | 138.52 | AD284333 |
| *Tecticornia halocnemoides* | 1.35 | 00/12/2017 | Australia | South Australia St. Kilda | -34.60 | 138.41 | AD284315 |
| *Frankenia pauciflora* | 2.51 | 00/12/2017 | Australia | South Australia St. Kilda | -34.73 | 138.52 | AD284313 |

**Table S3. Summary of raw, filtered and mapped reads for the sensitivity assessment (Section 3.1 of main text).**

| | | Sample conc ng/µl | Total number of reads | Reads after filtering | Reads mapped before removing PCR duplicates | Reads mapped after removing PCR duplicates |
|---|---|---|---|---|---|---|
| **restricted** | **Rep1** | 1 | 22629980 | 17290085 | 9029998 | 1077951 |
| | | 0.1 | 4974838 | 3832898 | 1993029 | 274094 |
| | | 0.01 | 472482 | 723012 | 313186 | 41729 |
| | | 0.001 | 4480408 | 7709131 | 896009 | 8712 |
| | | 0.0001 | 98207 | 174376 | 4807 | 915 |
| | **Rep2** | 1 | 42488364 | 32851172 | 17478695 | 1773158 |
| | | 0.1 | 1982450 | 1526600 | 750641 | 94012 |
| | | 0.01 | 4053066 | 274074 | 5463564 | 50076 |
| | | 0.001 | 1607459 | 2549181 | 1062429 | 19411 |
| | | 0.0001 | 87572 | 152861 | 2191 | 433 |
| | **Rep3** | 1 | 12708214 | 9893864 | 4955279 | 580195 |
| | | 0.1 | 3311566 | 2535517 | 1275934 | 164921 |
| | | 0.01 | 32902104 | 25163924 | 22090426 | 130102 |
| | | 0.001 | 1641021 | 2338448 | 1793728 | 15215 |
| | | 0.0001 | 76006 | 132990 | 2667 | 387 |
| **wider** | **Rep1** | 1 | 22629980 | 17290085 | 9042110 | 2798334 |
| | | 0.1 | 4974838 | 3832898 | 1996113 | 659723 |
| | | 0.01 | 472482 | 723012 | 313108 | 133033 |
| | | 0.001 | 4480408 | 7709131 | 890667 | 59449 |
| | | 0.0001 | 98207 | 174376 | 5044 | 1293 |
| | **Rep2** | 1 | 42488364 | 32851172 | 17502967 | 4741862 |
| | | 0.1 | 1982450 | 1526600 | 751435 | 242317 |
| | | 0.01 | 4053066 | 6274074 | 5470707 | 166030 |
| | | 0.001 | 1607459 | 2549181 | 1063092 | 73899 |
| | | 0.0001 | 87572 | 152861 | 2218 | 496 |
| | **Rep3** | 1 | 12708214 | 9893864 | 4961874 | 513990 |
| | | 0.1 | 3311566 | 2535517 | 1278140 | 414096 |
| | | 0.01 | 32902104 | 25163924 | 22117982 | 536016 |
| | | 0.001 | 1641021 | 2338448 | 1793428 | 105296 |
| | | 0.0001 | 76006 | 132990 | 3042 | 696 |

### 2.1 Statistical analysis output tables

**Table S4. Extraction optimisation- extraction kit.**

**A) Output from R version 3.5.1 (R Core Team 2018) where a one-way ANOVA was fitted to the data with extraction kit as a factor and DNA yield as the response variable.**

| | Df | Sum Sq | Mean Sq | F value | Pr(>F) |
|---|---|---|---|---|---|
| kit | 3 | 1.7567 | 0.5856 | 442.5 | **1.69e-05** |
| Residuals | 4 | 0.0053 | 0.0013 | | |

**B) Planned contrasts conducted between the different kits using the lsmeans package and the Tukey adjustment method.**

| contrast | estimate | SE | df | t.ratio | p.value |
|---|---|---|---|---|---|
| Combination - Stool | 0.036 | 0.036 | 4 | 0.98 | 0.77 |
| Combination - Plant | -0.16 | 0.036 | 4 | -4.5 | **0.036** |
| Combination - Soil | -1.11 | 0.036 | 4 | -30.54 | **<.0001** |
| Stool - Plant | -0.20 | 0.036 | 4 | -5.49 | **0.01** |
| Stool - Soil | -1.15 | 0.036 | 4 | -31.52 | **<.0001** |
| Plant - Soil | -0.95 | 0.036 | 4 | -26.028 | **0.0001** |

**Table S5. Extraction optimisation- recovery of plant DNA from soil**

**Output from R version 3.5.1 (R Core Team 2018) where a two-way ANOVA was fitted to the data with extraction kit and plant type as a factors and DNA yield as the response variable and testing whether there was an interactive effect between plant type and extraction kit.**

•

| | Df | Sum Sq | Mean Sq | F value | Pr(>F) |
|---|---|---|---|---|---|
| kit | 1 | 66.08 | 66.08 | 48.53 | **1.5e-05 ***** |
| plant | 2 | 10.29 | 5.14 | 3.78 | 0.053 . |
| kit:plant | 2 | 3.38 | 1.69 | 1.24 | 0.32 |
| Residuals | 12 | 16.34 | 1.36 | | |

**Table S6. Sensitivity assessment. A quasibinomial distributed Generalised Linear Model was fit to the data specifying species, reference and concentration as factors and including interaction terms. An ANOVA was then conducted on the model and the output is shown below**

| | Df | Deviance | Resid. Df | Resid. Dev | F | Pr(>F) |
|---|---|---|---|---|---|---|
| NULL | | | 89 | 79.621 | | |
| conc | 4 | 56.74 | 85 | 22.88 | 333.00 | **< 2.2e-16** |
| species | 2 | 17.85 | 83 | 5.026 | 210.19 | **< 2.2e-16** |
| reference | 1 | 0.002 | 82 | 5.024 | 0.036 | 0.85 |
| conc:species | 8 | 1.802 | 74 | 3.22 | 5.31 | **3.503e-05** |
| conc:reference | 4 | 0.017 | 70 | 3.21 | 0.10 | 0.98 |
| species:reference | 2 | 0.022 | 68 | 3.18 | 0.26 | 0.78 |

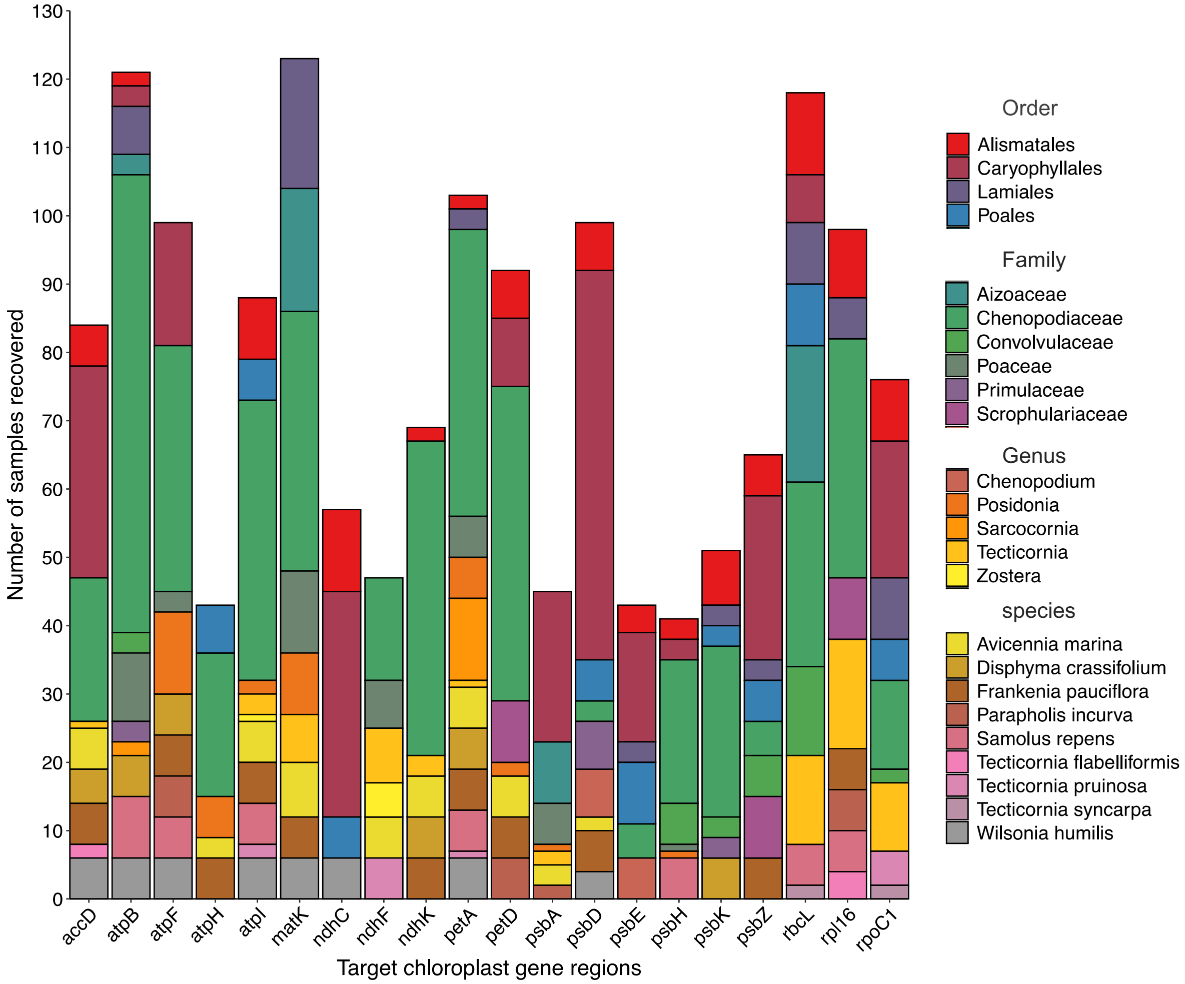


**Supplementary Figure 2. Species and gene recovery for the 10 species artificial DNA mixture for all taxa that were not put into the mixture, i.e., false positive detections.**
